# The Principle of Similitude in Biology: From Allometry to the Formulation of Dimensionally Homogenous ‘Laws’

**DOI:** 10.1101/250134

**Authors:** Andrés Escala

## Abstract

Meaningful laws of nature must be independent of the units employed to measure the variables. The principle of similitude (Rayleigh 1915) or dimensional homogeneity, states that only commensurable quantities (ones having the same dimension) may be compared, therefore, meaningful laws of nature must be homogeneous equations in their various units of measurement, a result which was formalized in the Π theorem (Vaschy 1892; Buckingham 1914). However, most relations in allometry do not satisfy this basic requirement, including the ‘3/4 Law’ (Kleiber 1932) that relates the basal metabolic rate and body mass, besides it is sometimes claimed to be the most fundamental biological rate (Brown et al. 2004) and the closest to a law in life sciences (Brown et al. 2004). Using the Π theorem, here we show that it is possible to construct an unique homogeneous equation for the metabolic rates, in agreement with data in the literature. We find that the variations in the dependence of the metabolic rates on body mass are secondary, coming from variations in the allometric dependence of the heart frequencies. This includes not only different classes of animals (mammals, birds, invertebrates) but also different aerobic conditions (basal and maximal). Our results demonstrate that most of the differences found in the allometric exponents (White et al. 2007) are due to compare incommensurable quantities and that our dimensionally homogenous formula, unify these differences into a single formulation. We discuss the ecological implications of this new formulation in the context of the Malthusian’s, Fenchel’s and Calder’s relations.

## 1. Introduction

The invariance of nature under scaling of units of measurement has been a powerful tool for the discovery of new phenomena, used for centuries in physics (Fourier 1822) and related areas (Macagno 1971). Being called dimensional analysis or principle of similitude (Rayleigh 1915), this methodology have been also applied to the solution of complex problems, ranging from the atmosphere behavior under a nuclear explosion (Taylor 1950) to airfoil prototypes (Bolster et al. 2011) and the formation of stars in galaxies (Escala 2015; Utreras et al. 2016)

Scaling invariance is desirable in any branch of science, not only restricted to physics, however, in many areas the fundamental relations have not yet been formulated in a form independent of the scaling of units. In the case of life sciences, particularly relevant are the allometric scaling laws, which relates a physiological variable with body size, which in almost all cases is measured by its mass. To fulfill the similitude principle implies that these relations can always be rearranged in terms of non-trivial dimensionless parameters/groups, however, allometric scalings almost never fulfill this property, which is also called dimensional homogeneity. Another way of formulating this issue, is as a phenomena that can be described with homogenous equations only with the aid of as many dimensional constants as there are variables (Bridgman 1922).

This problem is illustrated in the influential book ‘Scaling’ by Schmidt-Nielsen (1984), that shows dozens of relations between physiological variables in species and body mass, being the most notable one the so-called Kleiber’s Law, between basal metabolic rate (energy consumption per unit time) and body mass. None of the nontrivial relations satisfies the similitude principle, including the famous Kleiber’s Law, with one notable exception: the allometric relation between swimming speed of fishes and tail-beat frequency (Bainbridge 1958). When the fishes swimming speed is properly normalized dividing it by their body length, thus having frequency units as the tail-beats, the data among different fishes lies into a single curve in the same fashion as, for example, von Kármán (1957) unified different experiment of turbulent flows in pipes using only dimensional analysis.

The self-similarity displayed in Brainbridge’s allometric relation is appealing since it is the same displayed in physics and will be the main topic of this work. However, it is important to note first that there is another partial approach to guarantee similitude: fractal geometry. The possibility of having fractional dimensions allows to satisfy similitude with the use of an extra parameter, the Haussdorf or fractal dimension D. If the fractal dimension is properly chosen, is still possible to fulfill similitude for power laws with any fractional exponent.

In terms of dimensional analysis, fractal curves can be identified as incomplete similarities and such approach is the same used in renormalization group analysis in modern theoretical physics, since it can be proven the equivalence of incomplete similarity and invariance with respect to the renormalization group (Barenblatt 2003). Fractals belongs to the group of self-similar solutions of the second kind, in which the requirements of homogeneity and self-similarity are necessary to be satisfied only locally (Barenblatt 2003). On the other hand, self-similar solutions of the first kind (or complete similarities) that satisfies the more restrictive requirements of global homogeneity and self-similarity, can sometimes be obtained using the tools of dimensional analysis.

Fractals models of the allometric relations became popular in recent years, most notably since West et al (1997), that conjectured a universal ‘1/4 power law’ mass scaling for physiological variables and used fractal geometry to explain it from first principles. This gave considerable attention to fractal models for the allometric relations, however, also debate on the validity of such ‘1/4 power law’ universality (see for example the critical review by Hulbert 2014 or the comparison between theoretical predictions by Price et al. 2009). In fact, analysis of 127 interspecific allometric exponents discarded an universal metabolic relation (White et al. 2007) and using statistical analysis, Dodds et al. (2001) again discarded a simple scaling law for metabolic rates. Due to this intense debate, we will start analyzing possible fractals models, to then move towards complete similarity solutions using dimensional analysis tools (Π theorem).

This letter is organized as follows. We start studying the mathematical requirements to be a natural law in §2. We continue using dimensional analysis to study possible fractal models for the metabolic rates in §3. Section 4 continues with the search for a dimensionally homogenous metabolic rate equation and compare it against data taken from the literature. Finally in §5, we discuss the results and implications of this work.

## 2. Dimensional Constants and Homogeneous Equations

In the proper mathematical formulation of natural laws, a minimum requirement is to be expressed in a general form that remain true when the size of units is changed, therefore, meaningful laws of nature must be homogeneous equations in their various units of measurement, as was first observed by Fourier (1822). This requires, in addition to be a dimensionally homogeneous equation, that the constants with dimensions are restricted to only universal ones and to a minimum number, which in no case could exceed the total number of fundamental units of the problem.

However, in many areas (allometry being one of those) the phenomena can be described in homogenous equations only with the aid of as many dimensional constants as there are variables (Bridgman 1922). One example of such relations is the ‘3/4 Law’ (Kleiber 1932) that relates the basal metabolic rate (B) and body mass (M), described with an equation of the form

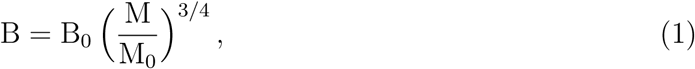

where B_0_ and M_0_ are normalization constants and therefore, in this relation each variable has its own constant with units. The ubiquity of this formulation for expressing relations probably comes from the fact that the procedure of fitting a curve to a set of points, gives such constants with dimensions and then it is natural to express the relation found in such terms.

The limitations of this approach can be better understood analyzing an specific problem, like the following thought experiment: let’s suppose that we are able to perform accurate experiments about the gravitational acceleration suffered by an object of mass m at a distance h from the surface of a planet. If we perform these experiments in the Earth and if any kind of friction or dissipation is negligible, taking the same approach used in allometric relations we will find that the acceleration data can be fitted by a relation of the form a = a_0_(1 + h/h_0_)^‒2^.

If we then travel to take measurements to a different planet, Mars for example, to perform the same experiment we will be able to fit the same ‘acceleration law’, a = ã_0_(1 + h/h̃_0_)^‒2^, but with different normalization constants ã_0_ and h̃_0_. Same case will happen if we take data from Jupiter, Neptune, etc. This methodology is exactly the one of as many dimensional constants (a_0_ & h_0_’s) as variables (a & h’s) and is the same scenario found in allometric relations (Schmidt-Nielsen 1984). Moreover, in this ‘one constant per variable’ approach to study the ‘acceleration law’, we have also one law per planet studied as in allometric relations where we have one law for each different classes of animals, with well documented intra-specific variations for invertebrates versus vertebrates, endotherms vs ectotherms, running vs resting, etc (White et al. 2007).

It is important to compare now the just mentioned methodology, to the one that lead to the most general ‘acceleration law’ of this problem. Thanks to Newton we know the general answer, not only for the Earth, but for any given spherical planet of mass M and radius R, which is given by

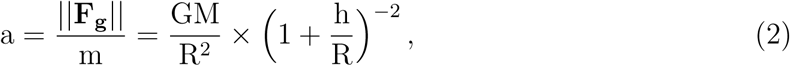

where G is the universal constant of gravitation. This solution comes from two well formulated laws, expressed by homogeneous equations in a form independent of the units of measurements: Newton’s Law of Universal Gravitation and his 2nd Law of motion. Such laws, were conjectured by Newton motivated on pure empirical observations and from them, a theory could be constructed that explain a whole variety of phenomena, in which the problem described by Eq. 2, is just one of many. In terms of dimensional analysis, this approach to the gravitational free-fall problem has 3 relevant dimensions (mass, length and time) and only one universal constant with units (G), compared to the previous case, in which as many constants as twice the number planets are needed to describe the same phenomena.

From the comparison of both approaches, the single well formulated dimensionally homogeneous solution (Eq. 2) and the one with as many normalization constants (a_0_ and h_0_’s) as variables (a and h’s), is clear that this multiplicity of laws and normalizations with units, are needed to explain variations of other relevant variables of the problem (R and M in this case). Also, it is straightforward to wonder if many of the exceptions in the Kleiber’s law and other allometric relations, are hiding variations of other relevant variables in the metabolic rate relation.

The difference between a dimensionally homogenous law that fulfill the similitude principle and phenomena described with the aid of as many dimensional constants as there are variables, is well known and used for centuries in physics. Newton was probably the first that realized this difference, when he conjectured his laws of motion and gravitation to explain pure empirical relations like the ones studied by Kepler, to describe the planetary motions around the Sun. Fourier was the first to realize the importance of units and that the laws must be homogeneous equations in their various forms. Maxwell established modern use of dimensional analysis in physics by distinguishing mass, length, and time as fundamental units, while referring to other units as derived. Contemporary physics allows a maximum of 3 fundamental constants (speed of light, Planck’s constant and G; Duff et al. 2002), which combined defines the natural base system to express the 3 fundamental units, called the Planck’s mass, length and time.

The rules required in the mathematical formulation of any natural law, was eventually formalized in the Π theorem discovered by Vaschy (1892), Buckingham (1914) and others. We will be use such theorem in the next sections, to search for a dimensionally homogenous law for the metabolic rates, starting with the less restrictive incomplete similarity solutions (fractal models) to then continue studying complete similarity ones.

## 3. Dimensional Analysis of Fractal Models

The Vaschy-Buckingham Π theorem defines the rules to be fulfilled by any meaningful relation aimed to be law of nature and it is a formalization of Rayleigh’s similitude principle. The theorem states that if there is a meaningful equation involving a certain number, n, of variables, and k is the number of relevant dimensions, then the original expression is equivalent to an equation involving a set of p = n – k dimensionless parameters constructed from the original variables. Mathematically speaking, if we have the following equation:

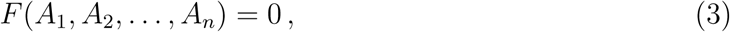

where the Ai are the n variables that are expressed in terms of k independent units, Eq 3 can be written as

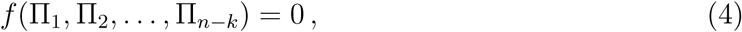

where the Π_i_ are dimensionless parameters constructed from the A_i_ by p = n – k dimensionless equations of the form 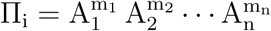.

We will follow the more general approach of Barenblatt & Monin (1983) to study a possible the fractal-like nature of biological systems, since it do not depend on a particular mechanistic model like the one in West et al. (1997) or assumes that the basal metabolic rate (V̇_O_2__) is directly proportional to the effective fractal surface of the body, as in West et al. (1999). We can construct a model by assuming that the metabolic rate, which is commonly measured in milliliters of O_2_ per minute and has dimensions [V̇_O_2__] = [M_O_2__]/[T], only depends on an absorbing capacity *β*_D_ with dimensions [V̇_O_2__][*L*]^‒D^, the body mass W with dimensions [M] and the density *ρ* with dimensions [*M*][*L*]^‒3^. The absorbing capacity *β*_D_ characterizes if oxygen-absorbing part of an organ could be approximated by a line (D=1), by a surface (D=2), by a volume (D=3)) or by an intermediate fractional dimension D (for more details, see Barenblatt & Monin 1983)

Because this model has n=4 variables (V̇_O_2__, *β*_D_, W & *ρ*) and k=3 independent units ([V̇_O_2__], [M], [L]), the Π theorem tells that one dimensionless parameter can be constructed, since n-k=1. To find the dimensionless parameter is straightforward by looking integer exponents such 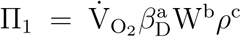 has no dimensions, which has the following solution: a = –1, b = – D/3 and c = D/3. Therefore, for this model it is possible to construct the following dimensionless quantity (Barenblatt & Monin 1983):

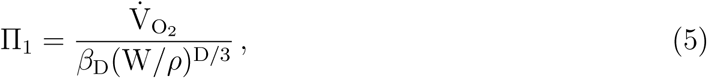

where it is straightforward to see that the allometric scaling for the oxygen consumption rate is V̇_O_2__ = AW^D/3^, with A = Π_1_*β*_D_*ρ*^‒D/3^. For a fractional dimension D=2.25, the usual 3/4 exponent of the Kleiber’s Law is recovered (Barenblatt & Monin 1983; Turcotte et al. 1998).

Another interesting quantity to look is the specific metabolic rate, V̇_O_2__/W, which from Eq. 5 takes the form:

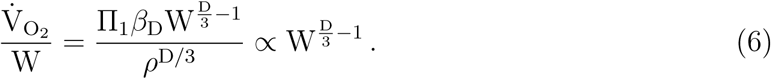

Since V̇_O_2__/W has dimensions of [M_O_2__][T]^‒1^ [M]^‒1^, it can be rearranged as equals to Π_1_*η*_O_2__*v*_o_, where *η*_O_2__ is an specific O_2_ absorption factor, with units [M_O_2__]/[M] and *v*_o_ that can be identified as a characteristic frequency, since it has inverse time units ([T]^‒1^). Assuming an absorption factor *η*_O_2__ independent of W, the characteristic frequency scales with body mass as 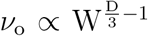, which can be directly compared against relevant rates like the respiratory and/or heart frequencies, in which there is considerable data on its scaling with W. This gives an independent Hausdorff dimension value needed to fulfill dimensional homogeneity.

For the case of mammals, heart frequencies scales as W^‒0.25^ (Brody 1945, Stahl 1967) which for a 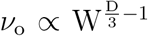, implies a fractional dimension of D=2.25, in agreement with the fractional dimension determined from the Kleiber’s Law. A similar result is obtained if we instead use, as characteristic frequency, the respiratory rates of mammals at rest (Calder 1968). In the case of sub groups, for example marsupials mammals have metabolic rates ∝ M^0.74^ and heart frequencies ∝ M^‒0.27^, which gives in both cases a dimension of D~2.2. In invertebrates such as spiders, the metabolic rates scales as M^0.59^ (Anderson 1970, 1974) and heart frequencies scales as M^‒0.41^ (Carrel & Heathcote 1976), quite different that in mammals, but both the metabolic rates and heart frequencies implies the same fractional dimension of D~1.8. In birds the scaling is again similar to mammals, proportional to M^0.72^ (Lasiewski & Dawson (1967) and M^‒0.23^ (Calder 1968), giving slightly different dimensions of D = 2.2 and 2.3 respectively.

We see a general consistency between the fractional dimensions D determined independently from the metabolic rates and heart/respiratory frequencies, regardless of the considerable variations of the D value among groups (especially in the case of invertebrates). It can be argued that such differences, are natural due to the different evolutionary stages among animal groups. Nevertheless, in the case of invertebrates is in addition required to argue in favor of oxygen-absorbing organs that are better approximated by fractal lines (D<2) than by fractal surfaces (2≤D≤3).

Another of the major issues in the exponent of the metabolic rate, is the change of the allometric scaling in the oxygen consumption under aerobic conditions (Weibel 2002), or maximal metabolic rate 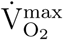. The scaling for 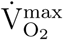 is approximately proportional to M^0.85^ (with slope variations ranging from 0.83 to 0.88; Savage et. al 2004, Taylor et al 1981, Dlugosz et al 2013, Weibel and Hoppeler 2005 and Bishop 1999) and the heart frequencies under such 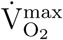 conditions, scales as M^‒0.15^ (Weibel and Hoppeler 2004, 2005). Again, both scalings gives the same fractional dimension, which in this case is D= 2.55. However, this kind of models based on fractal geometry requires now an aerobic change from D=2.25 to 2.55, in order to explain the change of allometric scaling under aerobic conditions. This assumption of time-dependent changes of the fractal network have not yet been observed, although it opens an interesting new possibility.

The requirement of adjustable fractal network under aerobic conditions is independent of the particular fractal model studied, unless the model assumes that the metabolic rate is directly proportional to the effective fractal surface of the animal, like in West et al. (1999), in which is not possible to explain the ∝ M^0.85^ scaling of the 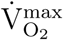. In such a case, the maximal possible scaling is ∝ M^0.75^, which correspond to volume filling surfaces (West et al. 1999).

## 4. A Complete Similarity Solution

In the different cases studied, we do not found a single and universal Hausdorff dimension D that explains allometric scaling for all organisms, but we found an excellent agreement in the dimension D determined from two different empirical allometric relations (metabolic rate and frequency). Nevertheless, the fractal model studied in §3 also assumes that the characteristic frequency *v*_o_ has all the D dependence on the W exponent, therefore in this particular model the variations in the W scaling of the V̇_O_2__ are secondary, coming directly from its dependence on *v*_o_. The agreement between the model and allometric relations, therefore suggests that indeed the W dependence in V̇_O_2__ and *v*_o_ might not be independent.

In such scenario, we are dealing with a similar case to the acceleration example studied in §2, where multiple laws and normalizations were needed to explain variations of other relevant variables of the free-fall problem (R and M). In the metabolic rate relation, the multiplicity of W scalings could be in fact needed to explain variations of another relevant variable: the characteristic frequency *v*_o_. This motivates us to explore *v*_o_ as an independent physiological variable that controls the metabolic rate V̇_O_2__.

Taking the characteristic frequency *v*_o_ as a variable in the metabolic rate, we will try now a different model with n=4 independent variables (V̇_O_2__, *η*_O_2__, W & *v*_o_) and which has now k=3 independent units ([M_O_2__], [M], [T]), therefore, from the Π theorem we know that n-k=1 dimensionless parameters can be constructed. The dimensionless parameter is again found by looking integer exponents such 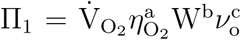 has no dimensions, which has the unique solution: a = b = c = –1. This implies that the dimensionless parameter is 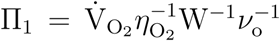 and if the metabolic rate relation depends only on this four variables, the Vaschy-Buckingham Π theorem states that it should be a function f such 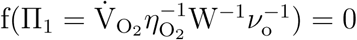 (Eq. 4). If function f have a zero that we called *ϵ*, such f(*ϵ*)=0, this implies:

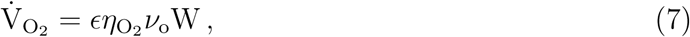

which is a self-similar solution of the first kind or complete similarity.

In order to confirm that the variations of the W scaling in the metabolic rate V̇_O_2__ comes from its dependence on *v*_o_, the self-similar solution found (Eq. 7) needs to be contrasted against empirical data. For that, it is required to check if the dimensionally homogenous relation has the slope of an identity, which must be unity within errors. For that purpose, we collected metabolic rates, masses and characteristic frequencies for different groups (mammals and birds) and aerobic conditions (basal and maximal).

Blue data points in Fig 1 are the basal metabolic rates V̇_O_2__ for mammals and the green ones, V̇_O_2__ for birds taken from Savage et al (2004) and Lasiewski & Dawson (1967) respectively. The respiration rates for both samples were taken from Calder (1968), converted to heart rates (f_H_) by multiplying a factor 4.5 for mammals and 9 for birds, values that were taken from the correlations between both rates listed in Schmidt-Nielsen (1984). The curves correspond to the least square fit to the data points, being 0.98 the slope of the V̇_O_2__ for mammals and 1.01 the slope of the V̇_O_2__ for birds. These slopes are both close to unity as expected to fulfill dimensional homogeneity. The normalization for birds slightly differs from mammals, however, this change in V̇_O_2__ normalization can be interpreted in our model as evidence in variations in the specific O_2_ absorption factor *η*_O_2__, since in Fig 1 we assumed *η*_O_2__ = 1[mlO_2_/g] for both birds and mammals. We choose the heart rate as characteristic frequency (*v*_o_ = f_H_) instead of the respiration frequency, because otherwise it increases the normalization shift seen in Fig 1, which suggests that a formulation with *v*_o_ = f_H_ will require less parameters to reach to an unique relation.

**Fig. 1.**
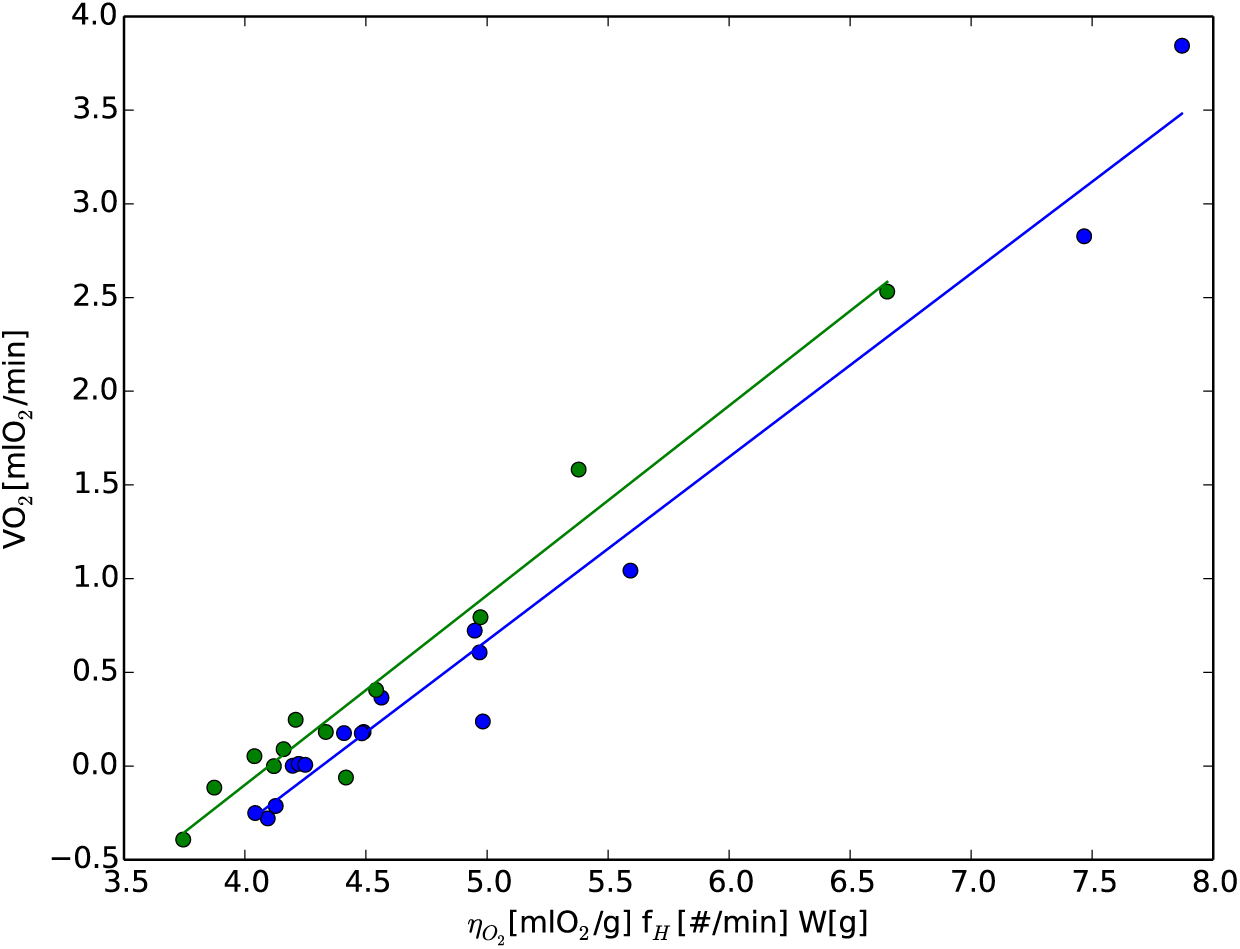
Basal metabolic rate V̇_O_2__ as a function of the heart frequencies and mass (with *η*_O_2__ = 1[mlO_2_/g]). The blue points correspond to mammals and green ones to birds data. The curves correspond to the least square fit to the data points, being 0.98 the slope or mammals and 1.01 the slope for birds.

Another major issue in the metabolic rate relation is the change in slope of the allometric W scaling when an animal exercises, in the change from V̇_O_2__ to 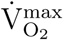, as mentioned earlier in the text. The red points in Fig. 2 display the V̇_O_2__ in a sample of resting mammals and the yellow ones are the corresponding 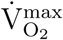 for the same species under maximal exercise. The sample of mammals was taken from Weibel & Hoppeler (2004), a group that has measured maximal and resting heart frequencies, with the corresponding maximal and basal metabolic rates taken mainly from Weibel, Bacigalupe et al (2004) and complemented with other references (Hinds et al 1993; Weibel 2000; Roef et al. 2002; Savage et al 2004; White et al 2006). The curves correspond to the least square fit to the data points, being 0.94 the slope of the V̇_O_2__ and 1.06 the slope of the 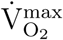.

**Fig. 2.**
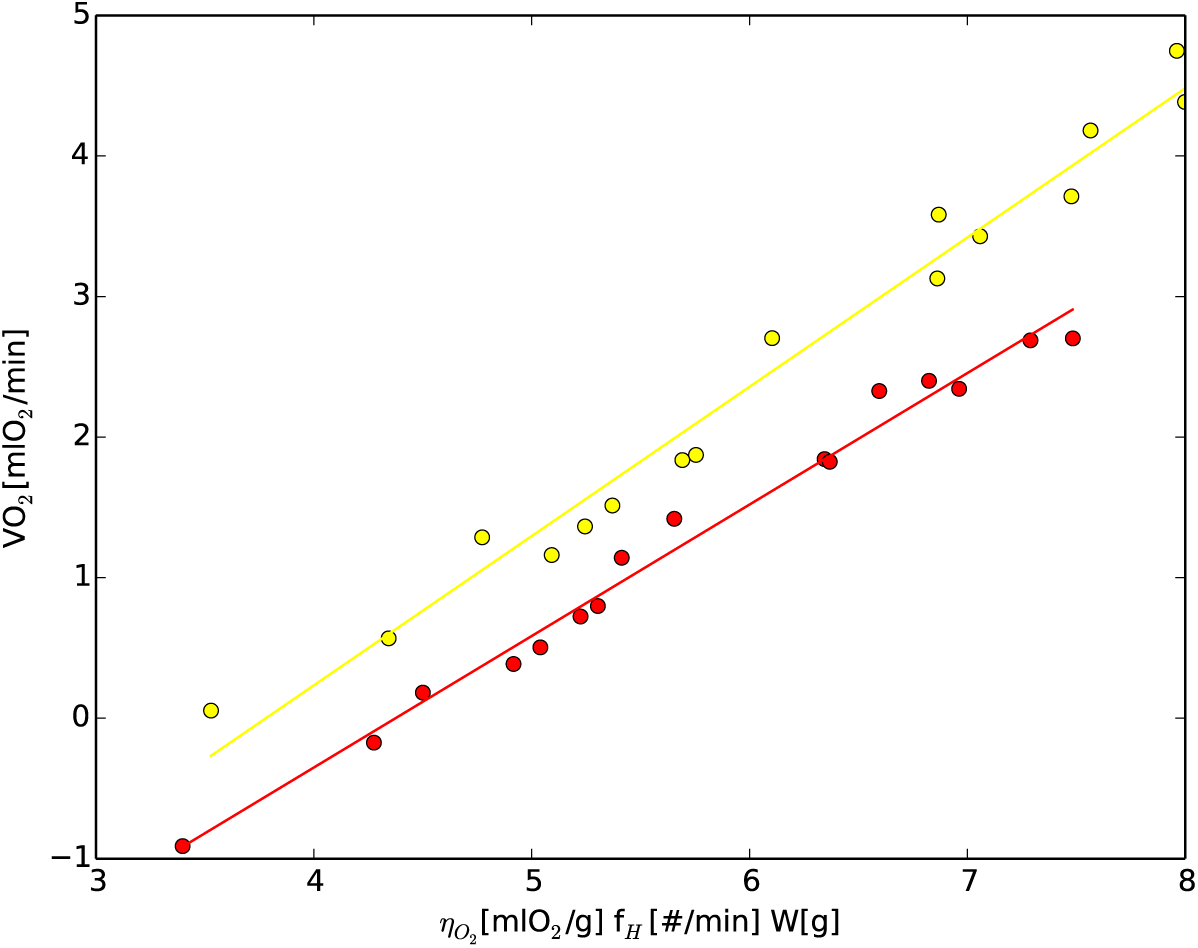
Same as Figure 1, but the yellow points correspond to maximal metabolic rates 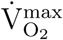 and the red ones are the corresponding V̇_O_2__ for the same mammals under basal conditions. The curves correspond to the least square fit to the data points, being 0.94 the slope of the V̇_O_2__ and 1.06 the slope of the 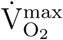

The larger deviation from slope unity can be explained in terms of the individual slopes of the sample of metabolic rates and frequencies showed in Fig. 2, in which V̇_O_2__ scales as W^0.69^ and the frequency as f_H_ ∝ W^‒0.26^, giving an V̇_O_2__/f_H_ proportional to W^0.95^. Similarly, the 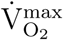 scales as W^0.90^ and 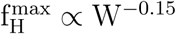 that gives an 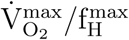 proportional to W^1.05^ instead of proportional to W^1.0^, as expected from Eq. 7 for both running and resting conditions to fulfill dimensional homogeneity. These variations from slope unity in the running and resting slopes are therefore due to the lower number statistics, because represent only a subsample of animals that has the three variables measured, in which the metabolic rates allometric exponents differs from the most accepted values (V̇_O_2__ ∝ W^0.75^ and 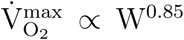). On the other hand, this support the multiplication/division of averages slopes used in §3, since the observed ones in Fig. 2 are almost the same slopes expected from both V̇_O_2__/f_H_ and 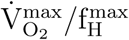. This further support the role of the heart frequency, f_H_, as an independent physiological variable that controls the metabolic rate.

The shift in normalization seen in Fig 2, can be interpreted as a change in the O_2_ absorption factor per unit mass, *η*_O_2__ in Eq. 7, which might be expected due to the shift in the animal’s internal oxygen demands between rest and exercise (Darveau et al. 2002; Weibel & Hoppeler 2005). Under basal conditions, the O_2_ consumption in celular respiration is mainly determined by the energy demands of basic maintenance processes in the tissues, compared to when an animal exercises that muscle work places a much larger O_2_ demand for energy supply (Weibel 2002). This change in the basic processes and organs that controls the energy demands of animals, should therefore imply a shift in the overall *η*_O_2__ of them, which in Fig 2 corresponds approximately to change in a factor of 5.

In order to quantify the change of *η*_O_2__ from basal to maximal exercise conditions, we estimate the O_2_ absorption factor per unit mass for the species under maximal exercise. Since only the product of *ϵ* × *η*_O_2__ can be extracted from the figures, we will focus on the relative changes by assuming *η*_O_2__ = 1 [mlO_2_/g] for resting animals (blue and green points in Fig 1 and red ones in Fig 2) and then search for the 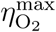 (yellow points in Fig 2) that minimizes the scatter in the relation. An absorption factor per unit mass of 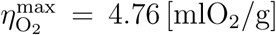 minimizes the scatter of the overall sample, being in such case 0.22 dex with respect to the relation:

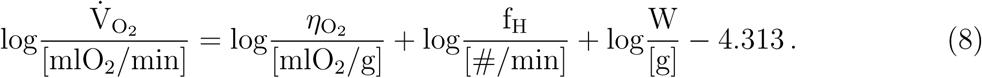

Fig 3 displays all the data in Fig 1 and Fig 2 with black points, with the just mentioned two different O_2_ absorption factor, namely, *η*_O_2__ = 1 [mlO_2_/g] for resting animals and 4.76 [mlO_2_/g] for those under maximal exercise. Also, the x-axes is re-normalized using the constant taken from the best fitted model (Eq. 8). This leads to a single relation consistent with a slope unity, being an identity relation like the black dashed line displayed in Fig 3.

**Fig. 3.**
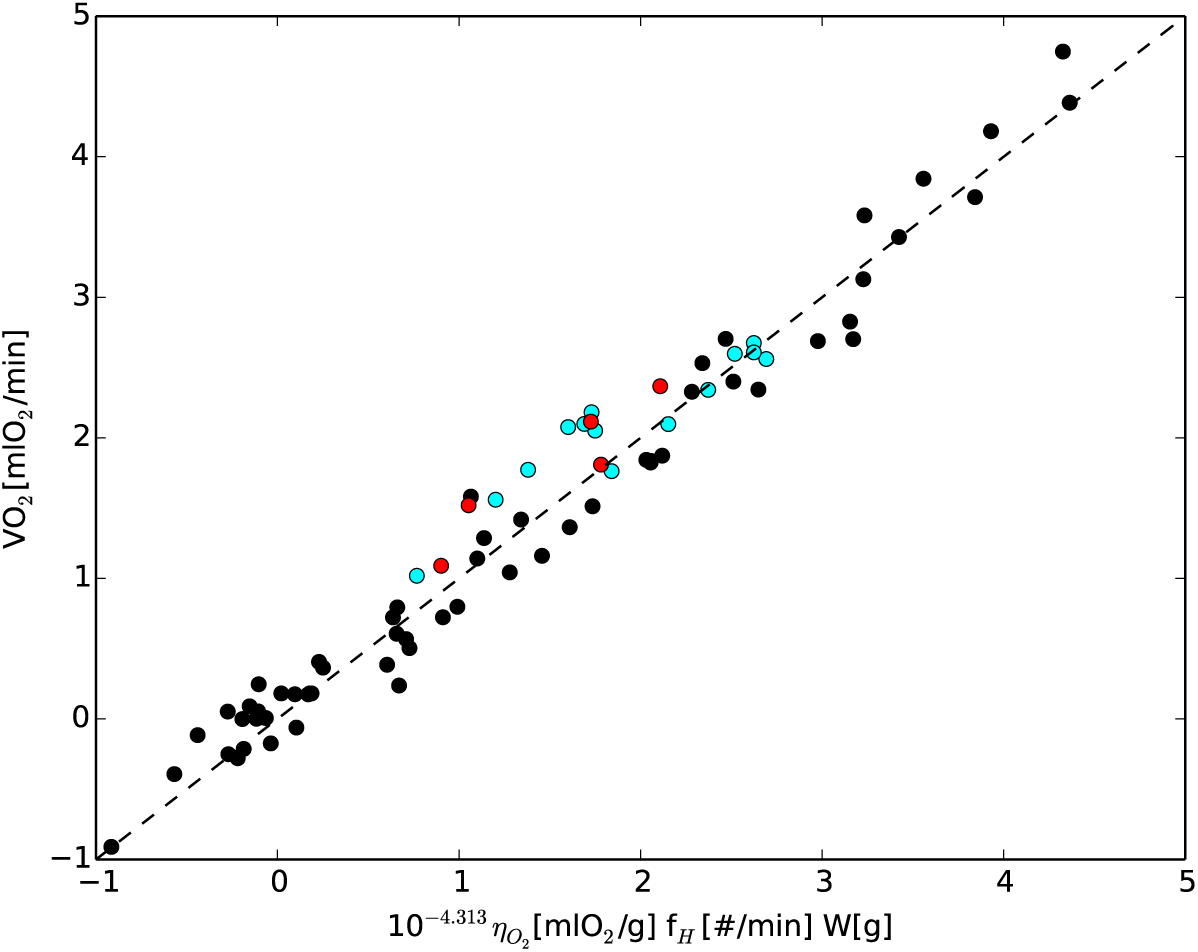
Black points are the same data showed in Figure 1 and 2, but we now assume *η*_O_2__ = 1 [mlO_2_/g] for resting animals and *η*_O_2__ = 4.76 [mlO_2_/g] for those under maximal exercise. The red points correspond to flying birds and the cyan ones to penguins under resting and running conditions, being both subsamples within the scatter of the relation (black points). The black dashed line denotes the identity between x and y-axes.

To check the relation found in Fig 3, with a different *η*_O_2__ for resting and running conditions, we also plotted data for two additional classes of animals. In cyan, we plot data for penguins under resting and running conditions, taken from Green et al 2005, and for flying birds in red (Berger, Hart & Roy 1970; Aulie 1971; Norberg 1996). Both subsamples, in which we again use *η*_O_2__ = 1 [mlO_2_/g] for resting subsamples and *η*_O_2__ = 4.76 [mlO_2_/g] for aerobic ones, are well described by the best fitted model given by Eq. 8, since both the cyan and red points lie within the scatter of the relation (black points).

For the introduced parameter *η*_O_2__ is hard to find direct measurements in the literature. Nevertheless, one possible estimation for *η*_O_2__ can be taken from the arteriovenous oxygen content difference, C_AO_2__-C_VO_2__, since it plays a role analogous to *η*_O_2__ in the Fick’s formula. Assuming that the cardiac stroke volume is directly proportional to its body mass W (Bishop & Butler 1995), it is possible to recover from the Fick’s formula an equation of the form of Eq. 7, but with a normalization dependent on C_AO_2__-C_VO_2__ instead of *η*_O_2__.

Factors of two difference in C_AO_2__-C_VO_2__ are in fact reported between resting and exercising/flying birds (*Columba Livia*) in Bishop & Butler (1995), between non-athletic and athletic mammals (Weibel & Hoppeler 2005) and even in the case of humans, only 3-months of moderate physical training increases the maximal oxygen intake 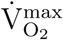 by 22.5%, mostly due to a change in C_AO_2__-C_VO_2__ (Detry et al. 1971). Taking these intra-specific differences into account, an overall factor of 5 change in *η*_O_2__ for different classes of animals could be even modest, specially noting that some species are intrinsically more aerobically evolved than others.

## 5. Discussion and Outlook

We showed that although in several cases the mass scaling in the metabolic rate and frequency differs from the predicted by the ‘universal 1/4 scaling’, their slopes are consistent with the same Hausdorff dimension D, if it is interpreted in terms of fractal geometry. This motivate us to propose unique homogeneous equation for the metabolic rates that includes f_H_ as independent controlling variable, for different classes of animals and for both resting and exercising conditions, which is in agreement with the empirical data. Our results demonstrate that most of the differences found in the allometric exponents (White et al. 2007) are due to compare incommensurable quantities, because the variations in the dependence of the metabolic rates on body mass are secondary, coming from variations in the allometric dependence of the heart frequencies on W. Therefore, f_H_ can be seen as a new independent physiological variable that controls the metabolic rates in addition to the body mass W.

The methodology presented in this work could have impact not only on the field of allometry, but also in life sciences in general. For example, the geometry of body cooling that is part of the debate in theories of metabolic allometry from even before Kleiber’s result, with our new formulation has no role, thus narrowing the discussion to the internal respiratory (O_2_) transport system. We consider that our result illustrates how powerful is to apply the Π theorem in the formulation of empirical biological laws, something that could also help to validate or discard theories in related areas such as ecology.

An open question is why the characteristic frequency changes its W scaling among different classes of living organisms and how this depends on the O_2_ transport problem. The reason for this might still has its origin on a ‘fractal like’ transport network, however, the dynamic scaling changes for 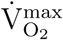 should also be partially related to changes in the energy demands (Darveau et al. 2002; Weibel 2002). Since O_2_ is a crucial ingredient in the cellular respiration and energy storage via ATP formation, how this is distributed along the body and where ends up being consumed, must both play a role in the integrated energy consumption of any organism.

A simple separation of both effects in terms of the complete similarity model presented here, might be to associate the change in the allometric scaling of *v*_o_ to a change in the transport network itself (which might be fractal as suggested by West et al 1997 and others), and the shift in normalization (*η*_O_2__) to the change of the organs that dominates the energy demand (from those associated to basic maintenance processes to those in muscle work as in Darveau et al. 2002).

The evidence that cells themselves have constant metabolic rates under ‘in vitro’ conditions (Gauthier et al. 1990), compared to the basal metabolic rates per cell that scales as W^‒0.25^ under ‘in vivo’ conditions (for a fixed cell mass; West et al. 2002), can be interpreted as the allommetric scaling of *v*_o_ only be related to the transport system. Moreover, even the case that dynamic changes in W scaling for 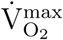 are due to changes on the organs that controls the energy demands of animals, this also implies a change in the transport network itself: from one reaching the organs of basic maintenance processes in the tissues, to one reaching where muscle work places, which in principle should have a different W scaling for *v*_o_.

An interesting limiting case for the *v*_o_ allometric scaling are plants, since they have metabolic rates linearly proportional to W (Reich et al. 2006), which in our model implies a characteristic (respiration) frequency *v*_o_ independent of mass. One simple interpretation is since most plants have relatively few living cells that carry out cellular respiration outside of their surface, they do not suffer the transport requirement and therefore such cells are in a condition more similar to the ‘in vitro’ one (Gauthier et al. 1990), having a characteristic frequency independent of the body mass.

Besides the successes of the relation found, other possible deviations are still documented in the literature. One particularly relevant is related to the temperature dependence, which is more obvious for Ectothermic systems since their body temperatures strongly varies with the environmental one. Gillooly et al. (2001) proposed a temperature corrected normalization for the Kleiber’s metabolic rate relation, based in the Arrhenius empirical formula for the temperature dependence of chemical reaction rates.

However, the temperature corrected metabolic relation still has residual variations around of factors 4-5 on the normalization of the rates between endotherms and ectotherms (Gillooly et al. 2001; Brown et al 2004). This is approximately the same variation in heart rates reported between endotherms (mammals and birds) and ectotherms (fish, anphibians, reptiles) in Lillywhite et al. (1999), suggesting that Eq 7 with the normalization corrected by an Arrhenius-type exponential dependence on temperature T, to take into account that chemical reactions proceed faster at higher temperatures, might be enough to also accurately account for the metabolic rates of ectotherms.

In terms of dimensional analysis, this can be derived assuming two additional variables T and T_a_ that controls the metabolic rate, being T_a_ an activation temperature defined as the activation energy E_a_ divided by the Boltzmann constant k_B_ (T_a_ ≡ E_a_/k_B_). In such a case, we have now a model with n=6 independent variables (V̇_O_2__, *η*_O_2__, W, T, T_a_, *v*_o_) that has now only k=4 independent units ([M_O_2__], [M], [T], [Θ]), therefore, n-k=2 dimensionless parameters can be constructed (Π_1_ = V̇_O_2__/W*η*_O_2__*v*_o_, Π_2_ = T/T_a_). In this case, the Π theorem states that there is an equation f(Π_1_, Π_2_) = 0 and if the function f is regular and differentiable, we can use the implicit function theorem to advocate the existence of a function Π_1_ = *ϵ*(Π_2_). Unfortunately, the functional dependence of *ϵ* on the second dimensionless parameter Π_2_ cannot be determined by dimensional analysis, therefore assuming an exponential form inspired in the empirical Arrhenius formula, namely *ϵ*(Π_2_) = *ϵ*_0_ e^‒1/Π2^, we get

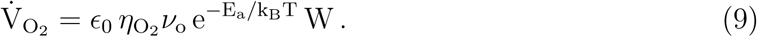

This equation can also be used to explain ecological phenomena and make predictions in a similar fashion as Gillooly et al. (2001) uses the temperature corrected relation on the metabolic theory of ecology (Brown et al 2004). To properly perform this, all the laws relevant to explain an ecological phenomena needs to be formulated in a form independent of the scaling of units (i.e. dimensionally homogenous equations in their various units of measurements). Once that is achieved, dimensional analysis can also be applied to solve ecological problems in a similar way as engineering does it with physical laws (Bridgman 1937).

The lack of well formulated laws in life sciences is probably the ultimate origin of the problems faced by theories such as the metabolic theory of ecology, which produces inaccurate statements (universal 1/4 power scaling; White et al 2007) and ecological implications (Duncan et al 2007). Well formulated empirical laws should precede theory, as happened historically in physics and chemistry. Also, only after their fundamental laws were formulated thru homogeneous equations in their various units of measurement and without exceptions, physics and chemistry reached the level of predictability characteristic of the exact sciences.

Nevertheless, is still possible to discuss implications of Eq. 9 for the few ecological relations formulated also in dimensionally homogenous equations. One of such is the population growth, which is an exponential controlled by the ‘Malthusian parameter’ or intrinsic rate of natural increase, r_m_, which has inverse time units and therefore should be associated to a frequency. Since its allometic scaling is r_m_ ∝ W^‒0.25^ (Fenchel 1974), is natural to associate it in our model with the characteristic (heart) frequency under basal conditions (*v*_o_ ∝ W^‒0.25^). However, less obvious is to find a causal connection between two frequencies that represents very different processes and timescales (internal circulation versus population growth). A possible link is in the total number of heartbeats in a lifetime N, which is approximately constant and equal to a billion for different mammals (Cook et al. 2006), then if the lifetimes scale inversely to the ‘Malthusian parameter’, we have: r_m_ ∝ 1/t_life_ ~ *v*_o_/N ∝ W^‒0.25^. Animal lifetime is clearly a more natural timescale for controlling the exponential population growth, which also has the same allometic scaling of population cycles (Calder 1983).

A constant total number N of heartbeats in a lifetime for mammals, t_life_ = N/*v*_o_, can also be used to relate the normalization 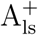 in the relation for the total energy consumed in a lifespan (Atanasov 2006), to our normalization (*ϵ η*_O_2__ in Eq 7). For N equals to a billion and converting 1 ltr O_2_ = 20.1kJ (Schmidt-Nielsen 1984), the 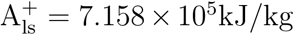 determined by Atanasov (2006) implies an 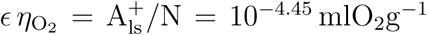. This is about the same number that can be determined independently from Fig 3, where the normalization of the relation is *ϵ η*_O_2__ = 10^‒4.313^mlO_2_g^‒1^ (noting that *η*_O_2__ = 1 [mlO_2_/g] for resting samples like those in Atanasov 2006).

Finally, for further fine-tuning of the metabolic rate relation, a coherent dataset of measurements is still required. Ideally, this needs to be for all the variables measured on the same animals and the original formulation of the metabolic law should stay strictly with measured quantities, for example with metabolic rates in ml O_2_ per min instead of energy-related units (watts or ergs/sec), to avoid assumptions in the conversion factors that produces undesirable extra scatter in the relation. Also, it will be interesting to test its predictions, for example to look for the temporal validity of Eq. 9 as an animal starts exercising and increases its O_2_ consumption. In ecology, an interesting test is to see if (Malthusian) population ecology parameters scale as v_o_ instead of universal 1/4, being a good candidate to see changes in allometic scaling of r_m_ of big outliers from ‘1/4 scaling’ in the metabolic relation under basal conditions, such as spiders (Anderson 1970, 1974) or other organisms (White et al 2007).

I thank Leonardo Bacigalupe for his help with the heart rates presented in Weibel & Hoppeler (2004) and the corresponding metabolic rates in Weibel, Bacigalupe et al (2004). I also acknowledge partial support from the Center of Excellence in Astrophysics and Associated Technologies (PFB 06).

